# Scale-free correlations and criticality in an experimental model of brain cancer

**DOI:** 10.1101/2022.07.10.499474

**Authors:** Kevin B. Wood, Andrea Comba, Sebastien Motsch, Tomás S. Grigera, Pedro Lowenstein

## Abstract

Collective behavior spans several orders of magnitudes of biological organization, ranging from cell colonies, to flocks of birds, to herds of wildebeests. In this work, we investigate collective motion of glioblastoma cells in an *ex-vivo* experimental model of malignant brain tumors. Using time-resolved tracking of individual glioma cells, we observed collective motion characterized by weak polarization in the (directional) velocities of single cells, with fluctuations correlated over many cell lengths. The correlation length of these fluctuations scales approximately linearly with the total population size, and these scale-free correlations suggest that the system is poised near a critical point. To further investigate the source of this scale-free behavior, we used a data-driven maximum entropy model to estimate the effective length scale (*n*_*c*_) and strength (*J*) of local interactions between tumor cells. The model captures statistical features of the experimental data, including the shape of the velocity distributions and the existence of long range correlations, and suggests that *n*_*c*_ and *J* vary substantially across different populations. However, the scale and strength of the interactions do not vary randomly, but instead occur on the boundary separating ordered and disordered motion, where the model exhibits classical signs of criticality, including divergences in generalized susceptibility and heat capacity. Our results suggest that brain tumor assemblies are poised near a critical point characterized by scale-free correlations in the absence of strong polarization.

## INTRODUCTION

Brain tumors are the most aggressive brain cancer, and remain the cancer with worst prognosis and shortest life expectancy. The standard of care treatment consists of resective surgery, radiotherapy and chemotherapy. Long term survival, nevertheless, has remained stagnant over the last thirty years in spite of major research efforts and multiple clinical trials [1–3]. Even when the tumor is resected, it always recurs, usually within a 1-2 cm margin of the original resection cavity. The tumor only rarely metastasizes to distant organs, but invades the surrounding normal brain, destroying normal brain areas, and disrupting brain function [4–6]. Brain cancer is the sole cancer that kills by direct invasion of surrounding normal brain tissue, rather than through metastasis of distal organs. Understanding the mechanisms of brain cancer growth and invasion are thus paramount to any future successful treatment of this disease.

Collective motion in brain tumors has been studied at the single cell and tissue level. However, our understanding of the mesoscale organization of brain tumor collective motion has not been studied in sufficient detail. In particular, collective motility requires close behavioural coordination between individual cells [7–10]. Such coordination relies on either direct cell to cell contact, close or indirect contact, or long-range information exchange between cells [11, 12]. The potential existence of such long-range communication might allow tumor cells to respond quickly to a number of insults, and thus provide robustness to tumor growth and progression. However, it is not clear whether such long-range correlations exist in brain tumors, and if so, how these large-scale patterns might arise from local interactions between nearby cells.

Collective motion arises in a large range of biological systems, from flocks of birds [13–18] to schools of fish [19, 20], from single cells [21–23] and insects [24–28] to populations of mammals [29–31] and has been intensely studied for decades in both natural systems [32–36] and robotics [37, 38]. Ordered motion could result from centralized, top-down mechanisms—for example, the presence of one or more “leader” cells that dictate the behavior of the others [39, 40]—or from emergent self-organization driven by local interactions between cells [33–35]. While these opposing modes of organization may lead to similar levels of order, the behavioral consequences of these two strategies are dramatically different. Self-organized systems are often characterized by a coherent response to external perturbations—for example, the presence of a predator in animal flocks—because local interactions facilitate an exquisite sensitivity to the changing environment, effectively propagating information from one region of the flock to another in a manner unseen in top-down organizational schemes [41–43]. Discriminating between these two modes of organization is challenging, and indeed they may often occur in combination. Our goal is to investigate the mode of ordered motion in brain tumors, and to do so, we draw on concepts from statistical physics, where emergent ordering is associated with well-defined features of simple mathematical models.

Scale free correlations are ubiquitous features of selforganized systems and have been observed in a number of biological systems [44, 45], including many exhibiting collective motion [13, 16, 17]. Correlations are considered scale-free when they lack a characteristic decay scale other than the size of the system. These correlations may enhance the system’s global response to perturbations by effectively linking the behavior of organisms across the entire population, even when direct physical interactions are limited to finite collections of local neighbors [13, 44]. Scale-free correlations can occur with or without strong ordering–that is, in the presence or absence of largely uniform behavior at the population scale. For example, in starlings, scale-free correlations in directional velocity occur in highly polarized flocks, where the distribution of velocity angles is extremely narrow [13, 15]. Similar scale-free correlations arise in physical systems with a continuous symmetry–in this case, the rotational invariance of the velocity angle–where low-energy fluctuations give rise to Goldstone modes such as long-wavelength spin waves [46]. By contrast, long-range correlations can arise in the absence of order–for example, swarming midges exhibit scale-free correlations despite show only weak levels of directional polarization. In physics, similar phenomena can occur when system parameters are tuned to a so-called critical point, where fluctuations become correlated across length scales and the system becomes exquisitely sensitive to small perturbations [46]. Recent work has argued for the existence of criticality in a wide range of biological systems [44, 47], including synchronization in neural ensembles [48, 49], brain activity [50, 51], long-range speed correlations in starling flocks [16], the dynamics of biochemical networks [52, 53] and protein folding [54], and the scale-free correlations in swarming midges [26, 55]. These two mechanisms of long-range correlations–those that occur with or without collective order–potentially serve different biological functions. For example, in flocks of birds, collective behavior may have evolved to promote lossless information flow throughout the group as they move through space, while swarms of midges might use collective behavior to stabilize their behavior against environmental perturbations, such as a predator.

In this work, we quantitatively characterize collective motion in an *ex-vivo* experimental model of malignant brain tumors using time-resolved tracking of individual glioma cells. We found that populations of glioma cells exhibit collective motion characterized by weak polarization in the (directional) velocities of single cells, with fluctuations characterized by correlation lengths that scale approximately linearly with the total population size. To investigate the source of this scale-free behavior, we used a data-driven maximum entropy model to estimate the effective length scale (*n*_*c*_) and strength (*J*) of local interactions between tumor cells. The model, which reduces to the classical XY model with nonlocal coupling, captures statistical features of the experimental data, including the shape of the directional velocity distributions and the existence of long range correlations. The length scale *n*_*c*_ and strength *J* of local interactions vary substantially across different populations, but they co-vary to fall on the boundary separating ordered and disordered motion, where the model exhibits classical signs of criticality, including divergences in generalized susceptibility and heat capacity. Our results suggest that brain tumor assemblies are poised near a critical point characterized by scale-free correlations in the absence of strong polarization.

## RESULTS

We analyzed glioma cells dynamics in a previously developed *ex-vivo*, explant model derived from the orthotopic implantation of genetically engineered NPA-GFP+ cells, in which we can track the location and velocity of fluorescently labeled individual glioma cells for up to 2 or 3 days (Materials and Methods). Briefly, this experimental model enables *de novo* the induction of glioma tumors trough the injection of different plasmids encoding 1) driver genes found in human gliomas and 2) genes encoding for luminescent and fluorescent markers in postnatal day 1 (P01) wild-type C57BL/6 mice. When animals become symptomatic, tumors are removed, and neurosphere cell cultures were established as described earlier [56–58]. Cells from these neurosphere cultures can then be implanted into adult C57BL/6 mice to reliably generate tumors (see Materials and Methods for details on the establishment of NPA tumors). NPA cells were then implanted into the brains of adult C57BL/6 mice [7, 56, 57]. Animals were euthanized at day 19 post-tumor implantation to generate explant brain tumor slices for time-lapse imaging on a laser scanning confocal microscope equipped with a tissue culture incubation chamber. We study a collection of 12 different glioblastoma populations drawn from 8 different explants–four from the inner tumor and four from the outer tumor region bordering the normal surrounding brain. We subdivided each explant into regions based on a previously developed classification scheme for glioblastoma cells [7]. While this scheme was originally developed to classify subpopulations of cells based on histological and statistical properties of the cell orientations, in the present context the subdivision scheme can be viewed as a box-like approximation for estimating finite-size scaling in the experimental system.

### Weakly ordered directional motion in glioblastoma populations

To quantify the population-level motion in the tumor, we estimated the position and velocity of each cell using semi-automated image analysis (Materials and Methods). At each time point, the population is described by a set of normalized (unit) velocity vectors {***s***_***i***_(*t*)} and a corresponding set of position vectors {***x***_***i***_(*t*)}, one for each cell *i* = 1, 2...*N*. To quantify the degree of global ordering in the population, we calculated the polarization *S*(*t*), which is defined as the magnitude of the population’s mean velocity; *S* = 1 when all cells move in the same direction, while *S* =0 if the velocities are uniformly distributed in all directions. We found that glioblastoma populations are weakly polarized–often exhibiting levels of polarization comparable to, or only slight exceeding, those in size-matched populations with velocities randomly drawn from a uniform distribution (compare black and red curves, Figure 1 and Figure S1). This weak polarization corresponds to broad but typically unimodal distributions of directional velocity (Figure 1, Figure S2). In contrast to starling flocks, which are highly polarized [13, 15], glioblastoma populations show minimal levels of polarization. Such weak polarization could be evidence that cells move largely independently, with little functional coupling between cells; on the other hand, weak polarization, alone, is insufficient to rule out collective behavior. Indeed, populations weakly ordered on the global scale have been shown to exhibit features of long-range collective behavior in a number of biological contexts, including swarming behavior of midges [17] and synchronous firing activity in neural populations [59, 60]. In these cases, fluctuations can be correlated over long spatial distances, even when ordering (i.e. polarization) is weak.

**FIG. 1.**
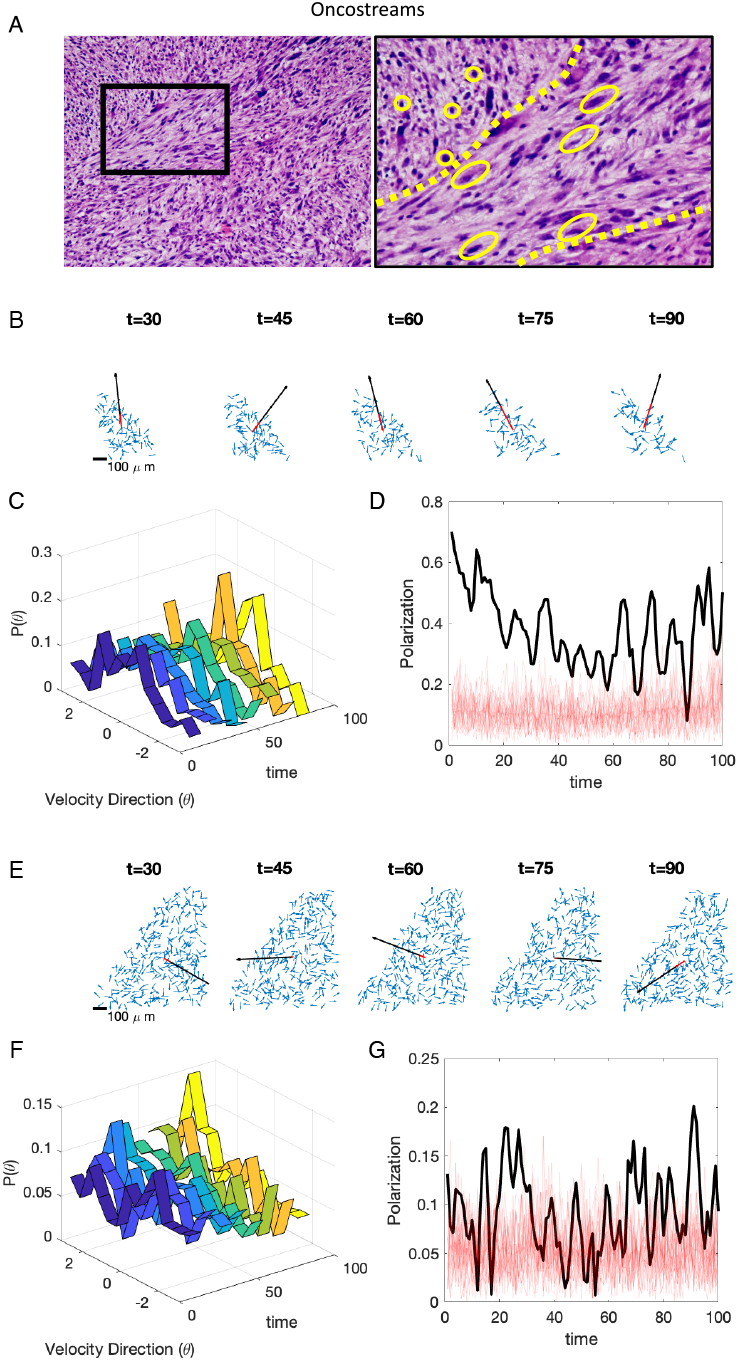
Cell velocities in glioblastoma populations exhibit weak directional order. A. Representative image of a 5um hematoxilin and eosin stained sections from an NPA mouse glioma tumor. The left image has been taken at lower power; and the black box is shown at higher power on the right. Oblong yellow outlines indicate elongated spindle-like cells within an oncostream –outlined by yellow stippled lines–while round yellow outlines indicate round cells located outside the oncostream. B. Snapshot of unit velocity vectors (blue) and average velocity vector (red) at different time points. Black arrow is a reference vector indicating full alignment (polarization = 1); length of red vector relative to black vector indicates polarization. C. Distribution of velocity directions over time (dark blue, early times; yellow, late times). D. Polarization over time for glioblastoma populations (black) and for 25 simulated datasets (size-matched to actual data) but with velocity directions drawn from a uniform distribution (light red; mean over all data sets in dark red). Polarization is defined by *S* = |⟨*s*⟩ |(1*/N*) Σ _*i*_ *s*_*i*_ |, where |*x*| is the length of vector *x* and angle brackets indicate an average over all cells. E-G: identical to B-D but for a second data set. See SI for all data sets.

### Glioblastoma populations exhibit scale-free correlations in velocity fluctuations

To determine if glioma populations exhibit correlated spatial fluctuations, we calculated correlations between directional velocity for cells separated by a given distance (*r*; in microns) for populations with different (time-averaged) spatial sizes (*L*). Correlation functions are defined by 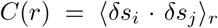, where 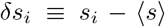 is the velocity of cell *i* in the moving reference frame where the population average velocity (⟨*s*⟩) has been subtracted out. Angle brackets ⟨⟩_*r*_ indicate an average taken over all cells separated by distance *r*.

We found that all populations exhibit weak local correlations over tens of microns (Figure 2, Figure S3). To characterize the decay of these correlations over space, we computed *r*_0_, (i.e. the point at which *C*(*r*) first crosses 0) for populations of different sizes. We found that *r*_0_ depends approximately linearly on the spatial size (*L*) of each population. This linear relation is expected to hold when the correlation length is much larger than the size of the system (i.e. when the correlated fluctuations are scale-free [61], and *r*_0_ can be interpreted as a correlation length scale; in contrast, when the correlation length is less than the system size, *r*_0_ grows as log *L*).

**FIG. 2.**
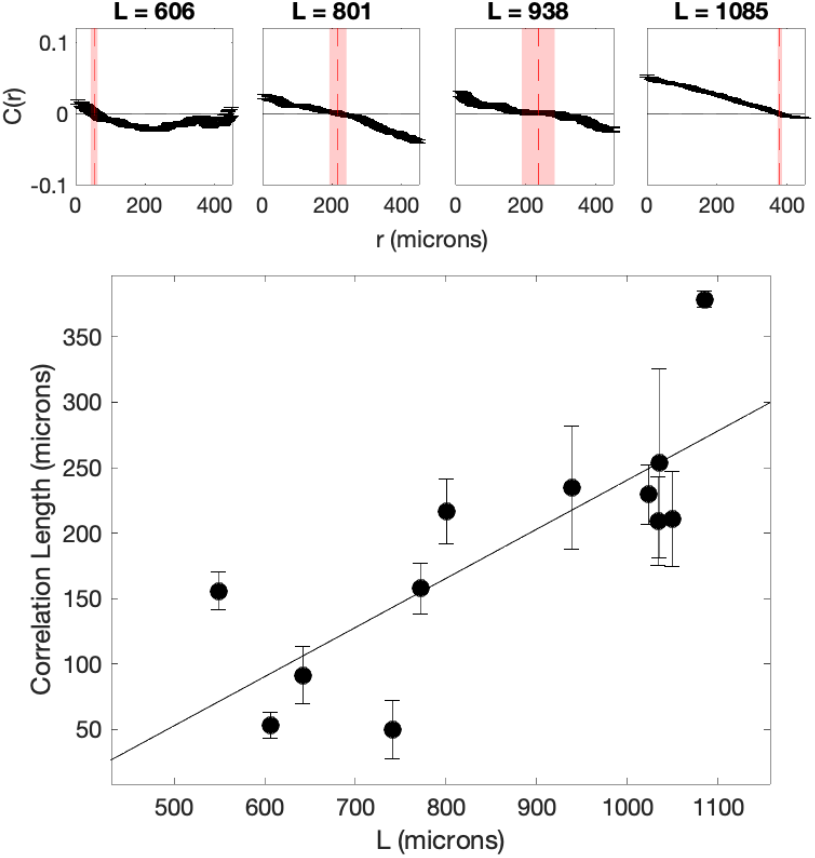
Velocity fluctuations are correlated over a length scale that depends on system size. Top panels: correlations between directional velocity for cells separated by a given distance (r; in microns) for populations with different (average) spatial sizes (*L*). Correlation functions are defined by 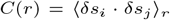, where 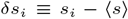 is the velocity of cell *i* in the moving reference frame where the population average velocity (⟨ *s*⟩) has been subtracted out. Angle brackets ⟨⟩_*r*_ indicate an average taken over all cells separated by distance *r*. Black markers: time-averaged correlations in a given population; red dashed line: estimated correlation length *ξ*, which corresponds to the crossover point *C*(*ξ*) = 0. Bottom panel: correlation length (*ξ*) for subpopulations of different sizes (black circles; error bars represent uncertainty in estimate of crossover point). Solid line is best-fit line (for visualization). See also Figure S3 for correlation functions of all populations.

In systems with a continuous symmetry, scale-free correlations can arise in the presence of global ordering (due to low energy Goldstone modes) or in the absence of global ordering (due to fine-tuning of the system to a critical point). In contrast to starling flocks, which exhibit strongly polarized velocities (*S >* 0.95), glioma populations exhibit weak polarization (typically *S* ≈0.3 or less) comparable to that observed in swarming midges. The observed scale free correlations–in the absence of strong polarization–provides strong evidence that glioma populations are poised near a critical point.

### A data-driven maximum entropy model captures statistical features of glioma migration

To develop a minimal effective model of glioma collective motion, we used the experimental velocity data to parameterize a simple maximum entropy model (Figure 3A). Maximum entropy methods have been widely applied to model biological phenomena [44, 62], including the collective firing activity of neurons [59, 60], the flocking behavior of birds [15, 16], and correlations in antibody diversity [15], drug interactions [63], or sequence motifs in biological polymers [64]. Maximum entropy approaches are closely connected to classical “inverse-problems” in statistical physics, which involve estimating (often unobservable) microscopic parameters from a set of macroscopic observables [65–69]. The maximum entropy approach allows one to incorporate a specific set of experimental observables into a model but, in a strict statistical sense, does not introduce additional structure. At its core, a maximum entropy model is consistent with a defined set of measurements but is otherwise as “unbiased” as possible. In the case of tumor cell populations, we would like to determine whether a minimal model that incorporates pairwise interactions of a fixed length scale is sufficient to reproduce the large-scale order observed in experiments. Following [15], we developed a maximum entropy model consistent with the local correlation structure measured in experiments—a quantity we refer to as *C*_int_. *C*_int_ is a single number that describes the average correlation of a cell’s velocity with that of its *n*_*c*_ closest neighbors; if cells tend to be directionally aligned with their neighbors, *C*_int_ approaches 1, while *C*_int_ is 0 if directional velocities are random.

**FIG. 3.**
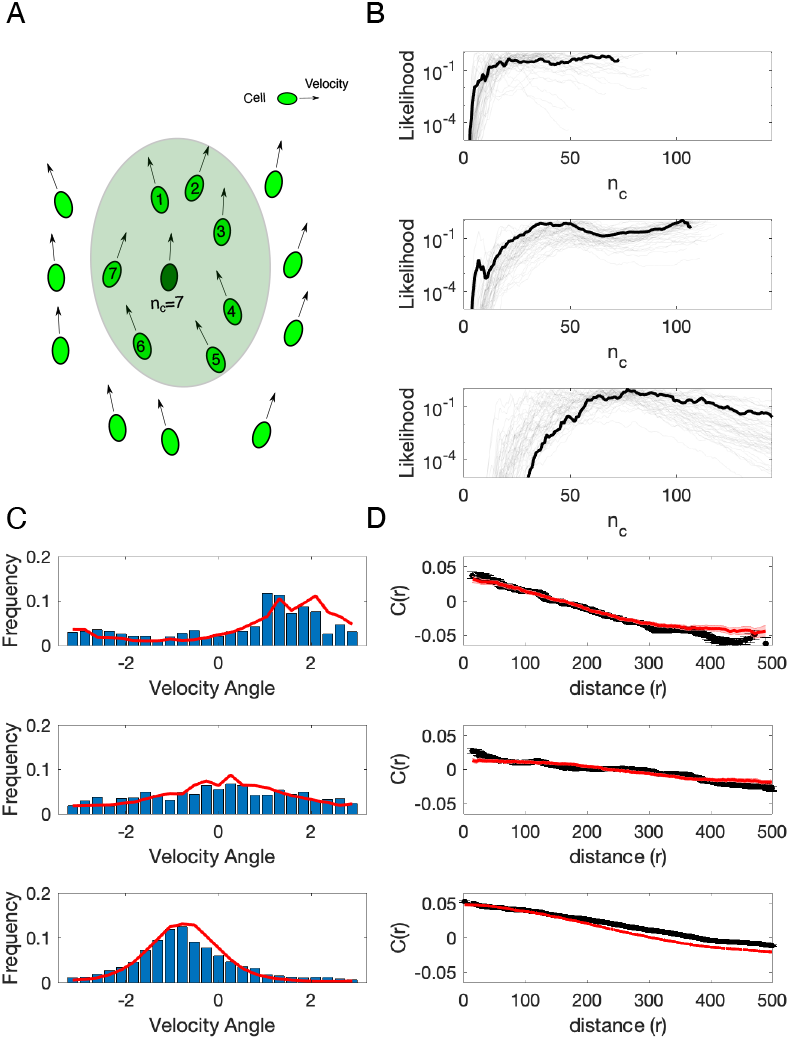
Maximum entropy model captures statistical features of glioblastoma populations. A. Moving cells (ovals, with arrows representing velocity) interact pairwise with the closest *n*_*c*_ neighbors in their local community. Shaded region indicates range of interaction for a single focal cell (dark green) for an illustrative value of *n*_*c*_ = 7. A coupling parameter *J* indicates the strength of interaction between pairs of cells, with larger *J* favoring stronger directional alignment. B. Likelihood (normalized) for the interaction range (*n*_*c*_) for three different populations; light curves show individual time points, dark curves are averages over time. C. Histograms of velocity direction (angle) for experiment (blue) and the maximum entropy model (red) for the same three populations as in panel B. D. Correlation functions calculated from experimental data (black) and for the maximum entropy model (red) for the same three populations as in panel B. See also Figure S4 and Figure S5 for full results on all populations.

The maximum entropy model consistent with *C*_int_ is formally identical to a non-locally coupled version of the XY model, which was originally developed in statistical physics to describe systems such as superconductors characterized by a continuous (planar) symmetry [46]. The model contains two free parameters: the length scale over which cells interact (*n*_*c*_), and the strength of those interactions (*J*). Our goal is to determine whether such a minimal model, containing only two scalar parameters, can reproduce measured features of the cell ensembles.

We estimated parameters (*J, n*_*c*_) for each population at each time point using a spin-wave approximation to calculate the partition function *Z* (see Materials and Methods). We note that while the spin wave approximation is strictly valid only in highly polarized systems, we found that the model with these parameter estimates qualitatively captures many features of our data. Different populations are characterized by widely variable (time-averaged) values of *n*_*c*_, with length scales ranging from tens of cells, in some cases, to hundreds of cells in others–nearly the size of the entire population (Figure 3B). To compare the model with experimental data, we used Monte Carlo simulations to estimate the distribution of velocity angles (Figure 3C) and correlation functions (Figure 3D) for the model with experimentally determined values of *J* and *n*_*c*_. Despite the simplicity of the model, it captures qualitative features of both the angular distribution and the correlation functions for nearly all populations without additional tuning of free parameters (Figure 3; see also Figure S4 and Figure S5).

Because the model suggests that cells are coupled over large length scales, with *n*_*c*_ sometimes approaching the size of the population, we also consider a mean-field (all-to-all coupled) version of the model. In this case, the model is characterized by a single parameter (*J*), and estimating *J* from experimental data reduces to a classic inverse problem in statistical physics. The advantage of this simpler model is that parameter estimation no longer relies on the spin-wave approximation, and instead, the maximum likelihood estimate of *J* can be calculated analytically in terms of the experimental observable *C*_int_ (Material and Methods). Somewhat surprisingly, the mean field model captures the angular distribution of velocities nearly as well as the full model Figure S6). As a numerical control, we confirmed that such agreement does not occur in randomly generated data sets of size-matched populations, confirming that the qualitative agreement between the model and experimental data is unlikely to arise from finite size statistical effects S7.

### Effective model of glioma dynamics is poised at a critical point

Glioma cells exhibit scale-free correlations in the absence of polarization, which provides evidence that the system is poised near a critical point. We therefore wondered whether the effective maximum entropy model is also poised near a critical point: that is, whether the experimentally derived parameters (*J, n*_*c*_) correspond to a critical point of a model. The absence of criticality in such a model may point to alternative explanations for the experimental observations.

To characterize the phase diagram for the maximum entropy model (i.e. an XY model with nonlocal coupling determined by *J* and *n*_*c*_), we used Monte Carlo simulations to generate representative velocity distributions for different values of *n*_*c*_ and *J*. For each parameter pair, we calculated the polarization order parameter (the mean polarization across trials), the generalized susceptibility *χ*, which corresponds (up to a proportionality constant) to the variance of the polarization, and the heat capacity, which measures fluctuations in the effective energy. We then estimated the critical surface in the (*J, n*_*c*_) plane to be the curve in parameter space that corresponds to a peak in *χ*. We also verified that the system exhibits sharply increasing polarization and a peak in the generalized heat capacity at the critical surface. For all simulations, cell positions (and therefore cell density) were taken from a representative experimental data set.

The phase diagram (Figure 4A) is divided by the critical surface into an ordered region (upper right) and disordered region (lower left, gray). To investigate whether the experimental populations are poised near the critical surface, we plotted the estimated parameter values (*J* and *n*_*c*_) for each population (triangles, squares, and circles) on the phase diagram. Indeed, we find that all populations are characterized by parameter pairs that lie close to the critical surface. Modulating the parameters–for example, by simulating a system with a fixed (experimentally determined) value of *n*_*c*_ but *J* values that differ from subcritical to supercritical (as indicated by the dashed line connecting points X and Y, Figure 4A)–indicates that the parameters estimated from experiments indeed occur near a critical point, where polarization begins to rapidly increase and *X* and the heat capacity peak (Figure 4B).

**FIG. 4.**
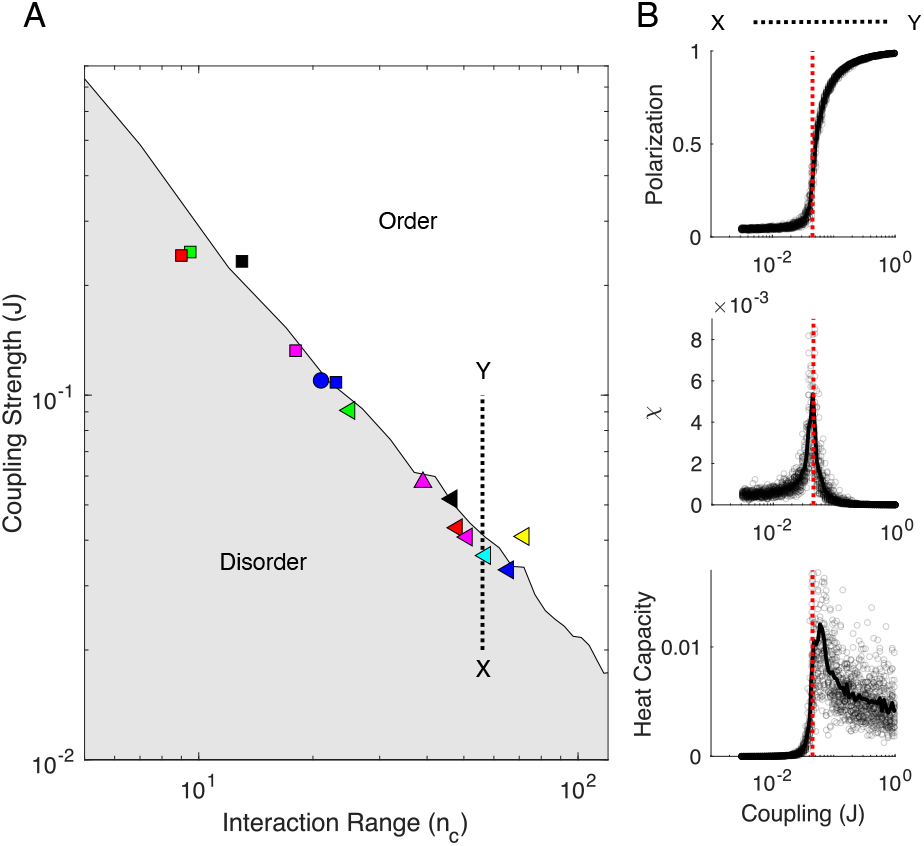
Experimental estimates of cell-cell coupling parameters are poised near critical point of non-locally coupled XY model. A. Phase diagram shows regions of disorder (gray) and order (white) for different values of the coupling strength (*J*) and interaction range (*n*_*c*_). Phase boundary was estimated from Monte Carlo simulations of the maximum entropy model (Materials and Methods). Markers (squares, triangles, circles) show estimates of *J* and *n*_*c*_ from different glioblastoma populations. Dashed black line shows an example trajectory through parameter space–in this case, from a point X in the disordered region to a point Y in the ordered region–that crosses the critical surface by increasing *J* at a fixed value of *n*_*c*_. B. Polarization (top), generalized susceptibility χ (middle), and generalized heat capacity (bottom) calculated from Monte Carlo simulations of the model as parameters are varied along the dashed trajectory connecting points X and Y in panel A. Gray circles represent Monte Carlo simulations across a range of coupling strengths (*J*) at fixed *n*_*c*_ = 58 (corresponding to the estimated *n*_*c*_ for one representative data set); solid lines are averages over a total of 100 simulations. Dashed red line indicates estimated value of coupling *J* for one particular data set (green triangle in panel A) that lies near the critical surface. Susceptibility (*x*) and heat capacity are defined as the variance of polarization and energy, respectively, across independent realizations.

We stress, however, that criticality in this model (alone) is supportive, yet insufficient on its own to infer criticality in the glioma populations, both because of technical limitations in distinguishing critical and subcritical systems (see SI) and, more importantly, because this effective model may not correspond to the true physical mechanisms underlying the motion. While the maximum entropy model accurately captures qualitative behavior of the glioma populations–and in doing so, provides a parsimonious effective model for the data–the strongest evidence for criticality is the (model-independent) observation of scale-free correlations in the absence of ordering.

## DISCUSSION

We’ve shown that collective motion in brain cancer populations is characterized by scale-free correlations in the absence of strong polarization, providing evidence that glioma cells may be poised near a critical point. The statistical features of this motion are largely captured by a simple, effective model which is, itself, also poised at a critical point.

Our work complements a growing number of studies that indicate biological systems may operate at or near criticality [16, 44, 45, 48–54, 70–72]. On the other hand, the behavior of glioma cells is substantially different than that of starling flocks, where scale-free correlations and strong polarization may be related to low-energy Goldstone modes that are ubiquitous in systems with similar rotational symmetry [15, 33]. Interestingly, starling flocks do show signs of criticality in the distribution of speed (as opposed to velocity direction) [16]. In the case of glioma cells, it is not clear if signatures of criticality are evident in the collective behavior of other properties, though it is an exciting avenue for future work.

It is important to keep in mind several limitations of our study, both technical and conceptual. First, the experiments are performed in an ex-vivo model, which allowed us to measure cell velocities and track movement for up to 72 hrs. As always, however, one should use caution when extrapolating from laboratory results to in vivo dynamics, though recent data have demonstrated similar dynamics using multiphoton intravital imaging of glioma [7]. In addition, our use of a maximum entropy approach comes with important caveats. While the model captures many statistical properties of the underlying experimental data, it should be viewed as an effective model, not necessarily a true physical representation of the underlying system. For example, the model suggests that pairwise interactions between cells are long-range–often extending across tens of cells or more–but there is no guarantee that the true biological interactions, whose mechanisms are, at this stage, largely unknown, extend over this scale, though for example cytokine secretion or recently discovered Ca++ mediated long-range intercellular connectivity in glioma might serve as potential biological substrates [73]. Instead, long-range interactions in the model could represent effective interactions that appear artificially long-range because (for example) the local cell dynamics have not reached equilibrium on the timescales of our measurements [74, 75], meaning that the effective interactions represent many fast physical interactions between different pairs of cells. Perhaps more importantly, the criticality observed in the data-driven model does not, alone, mean that the glioma populations are also at criticality. Similar effective models have been widely used over the last decade, but it is generally understood that features of these models–including criticality and the absence of higher-order interactions– do not necessarily reflect similar features in the underlying system [47, 76–81]. In this case, while the effective model accurately captures qualitative behavior of the glioma populations, the strongest evidence for criticality is the (model-independent) observation of scale-free correlations in the absence of ordering.

There are also a number of technical limitations that bear mention. We focused on a model with topological (rather than metric) connections between cells. For these populations, where the densities of cells are similar across populations, we expect that both approaches would yield similar results, though imposing a different coupling structure could improve the agreement between model and experiment [13, 17]. In addition, we have used free boundary conditions, in part because confining the boundary cells (using fixed boundary conditions) would, in some cases, significantly reduce the number of cells for analysis. Previous work in starling flocks showed that these boundary conditions can indeed make a difference in the inferred values of *J* and *n*_*c*_, though their product remains largely constant [15]. Correcting for these boundary conditions might provide a more accurate fit to, for example, simulated systems with known control parameters. In this case, however, our goal was not to precisely estimate experimental parameters, and because the model captures experimental results reasonably well, we have not further investigated these issues.

Our results raise a number of open questions for future work. From a physics perspective, more detailed physical models may shed additional light on quantitative features of glioma migration that are not captured by the simple model used here. For example, we recently showed that a model of collective motion that incorporates cell shape can produce a rich collection of phenomena that includes nematic ordering and qualitatively distinct types of disordered motion [10] similar to that seen in glioma populations [7].

Perhaps most interesting, our results imply the existence of (effective) long range communication within glioma tumors, but the physical basis for this communication is not known. Previous work indicates that natural insect swarms exhibit long-range correlations [26], but interestingly, these correlations only appear in response to different environmental perturbations in laboratory populations [28, 82]. Similarly, it may be possible to investigate these mechanisms in glioma populations using controlled laboratory conditions that strip away complexities of the ex-vivo tumor environment. From a bioloigical perspective, several mechanisms exist that could play a role in transmitting signals across large portions of glioma. For example, cytokines or neurotransmitters could be released by glioma cells, diffuse throughout the tumor tissue, and thus alter the behavior of distantly located cells. Equally, networks of microtubes connecting glioma cells have recently been described and shown to provide signals to other cells via Ca++ transients. So far, only shorter distance communication has been shown, but if microtubes are truly functional across larger distances, they could be an anatomo-physiological substrate of long range communication. Further, criticality could help understand and explore the recalcitrant robustness of these tumors, when exposed to therapeutic modalities such as X-rays, and chemotherapy. Criticality could play a role in glioma cells responding to resective surgery; for example, no matter how much tumor is resected, cells located within 1-2cm from the resection cavity eventually reconstitute the tumor. Thus, it might be possible that low density cells might need to grow to a certain size before long range communication could be use to support cell replication and especially cell invasion. Finally, a role for the recently discovered brain innervation of tumors could be to directly connect distant parts of the tumor, i.e., tumor cells could signal to innervating neurons which, through neural networks would then signal back to distant tumor regions. These are plausible mechanisms that could support long range communication in glioma, and are compatible with the time lengths described in our experiments.

## MATERIALS AND METHODS

### Time-lapse confocal imaging in explant brain glioma slice culture model

To analyze glioma cells dynamics, we used a brain tumor explant model for imaging ex-vivo. Glioma tumor cells were obtained from a genetically engineered mouse glioma model generated in our lab as previously described (Sleeping Beauty Transposase System) [56–58]. This model enables de novo generation of glioma tumors trough the injection of different plasmids encoding the genes of interest in postnatal day 1 (P01) wild-type C57BL/6 mice. The plasmid sequences used to generate these tumors were: 1) The transposase & luciferase enzyme expression (pT2C-LucPGK-SB100X), 2) the NRAS gene expression (pT2CAG -NRAS-G12V), 3) the short harpin for downregulation of p53 protein (pT2-shp53-GFP4) and 4) the short harping for ATRX (pT2-shATRx-GFP4). This glioma tumor cells named NPA exhibit the overexpression of the NRAS protein, the knockdown of p53 and the downregulation of ATRX. shp53 and shATRx plasmids also contain the gene for the expression of the enhanced green fluorescent protein (EGFP). These signaling pathways are altered in human gliomas. Cells were generated from genetically engineered glioma tumors and cultured in DMEM/F12 media supplemented with 2% B-27, 1% N-2, 1% of Penicillin-Streptomycin, 0.2% of Normocin™ and growth factors hFGF and hEGF at 20 ng/ml and maintained in a 37 C incubator supplied with 95% air and 5% CO2. Tumor were induced by intracranial implantation of 3 × 104 NPA cells (NRAGV12, shp53, shATRx) glioma tumor cells in C57BL6 mice. Mice were held in a pathogen free, humidity and temperature controlled vivarium on a 12:12 hour light:dark cycles with free access to food and water following. Prior to implantation, mice were anesthetized using an intra-peritoneal (i.p.) injection of the anesthetics Ketamine (120 mg/Kg) and Dexmedetomidine (0.5 mg/Kg). Following anesthesia, Carprofen (5.0 mg/Kg) was administered subcutaneously. The skull of the mouse was then immobilized in a stereotactic device. A hole was made using a 0.45 mm drill bit at coordinates corresponding to the striatum (0.5mm anterior and 2mm lateral right from the bregma). Cells were injected with a Hamilton syringe at a dorsoventral position of 3.5 mm into the striatum. After injection, the incision was sutured and immediately following surgery, the animals were recovered from anesthesia using Atipamezole via i.p. injection (1.0 mg/Kg) to reverse the Dexmedetomidine. A single subcutaneous injection of Buprenorphine (0.01 mg/Kg subcutaneous) was administered as post-operative pain relief. Sutures were removed 10 days after surgery [7, 56].

Tumors were allowed to grow for 19-21 days. At 19-21 days post-implantation animals were euthanized to generate the brain tumor slices explants for imaging. To generate brain tumor explants, brains were embedded in 4% low melting temperature agarose and kept on ice until solidification. Embedded brains were then immersed in ice-cold and oxygenated DMEM media without phenol red and then sectioned in a Leica VT100S vibratome (Leica, Buffalo Grove, IL). 300 mm thick brain tumor sections were transferred to laminin coated cell culture insert (Millipore Sigma, USA) placed into a 27mm diameter dish (Thermo Scientific) with D-MEM F-12 media supplemented with 25% FBS and Penicillin-Streptomycin. All steps were performed under sterile conditions in a BSL2 laminar flow hood. Tumor slices were then maintained in a cell culture incubator at 37 C with a 5% CO2 atmosphere. After 6-18 hours’ media was replaced with DMEM-F12 media supplemented with B27 2%, N2 1%, Normocin 0.2 %, Penicillin-Streptomycin 10.000 U/ML and growth factors EGF and FGF 20 ng/ml. After, tumor explants were transferred to the incubator chamber of a single photon laser scanning confocal microscope model LSM 880 (Carl Zeiss, Jena, Germany). For tumor imaging the incubation chamber of the microscope was maintained at 37°C and 5% CO2. Images were acquired in a time-lapse frame of ten minutes for 100-300 cycles. The movies were originally described in reference [7].

### Image Analysis

To track the evolution of the cells, we use the software *Fiji* with the plugin *TrackMate*. We use as parameters for the cell size (called ‘blob’) 20*μ*m and a threshold of 1 together with the DoG method (Difference of Gaussian detectors). Each experiment gives several paths denoted by **x**_*i*_(*t*) where *i* is an index for the cell and *t* represents the time. The paths are however erratic thus we apply a filter to smooth the trajectories over time (see figure S10). As a filter, we use a Gaussian kernel with standard deviation *σ*^2^ =2 and a stencil of 9 points:

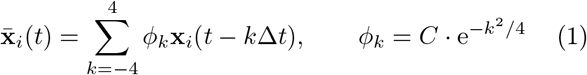

where 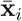 is the smooth trajectory, Δ*t* = 10 mn is the time step between successive image and *C* is such that 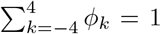. From the smooth trajectories 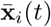, we then estimate the velocities of the cells 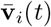 using a finite difference:

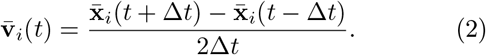

For each velocity vector 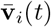, we estimate a corresponding velocity angle *θ* _*i*_(*t*) ∈ [0, 2 *π*) (see figure S10). For subsequent analysis, all velocity vectors are normalized and referred to as ***s***_***i***_

### Classification flock-stream-swarm

We subdivide glioma populations into connected subpopulations which we refer to as flocks, streams, or swarms (see figure S11-left). This empirical classification scheme was developed in previous work [7] to segment populations into subgroups with similar histological and statistical properties, as described briefly below. In the present context, these classifications are not particularly important, and instead one may view this as a systematic way of subdividing populations for finite-size scaling, which is necessary because the number of experimental populations is limited (see [83] for further discussion of these issues in limited data sets).

To classify in which category an experiment is (i.e. flock, stream or swarm), we analyze the collection of velocity angles { *θ* _*n*_}_*n*=1..*N*_ (where *N* is the sample size). We determine three densities for each pattern (see figure S11-right):

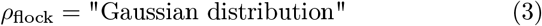

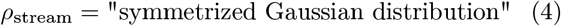

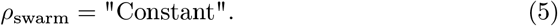

We then evaluate the likelihood of the velocity angles {*θ* _*n*_}_*n*=1..*N*_ given each distribution, for instance:

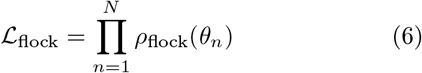

Finally, we compare the three likelihoods (i.e. ℒ _flock_, ℒ _stream_ and ℒ _swarm_) and select the pattern with the highest likelihood.

### Calculating correlation functions

The correlation function is given by 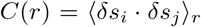, where 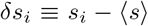 is the velocity of cell *i* in the moving reference frame where the population average velocity (⟨ *s*⟩) has been subtracted out. Angle brackets ⟨ ⟩ _*r*_ indicate an average taken over all cells separated by distance *r*. In practice, we calculated correlation functions by first calculating pairwise distances and dot products 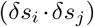 for all pairs of cells in a image at a given time. We repeated this process for all images in a time series and combined the data from all time points into a single pair of lists (distances and corresponding dot products, ordered according to distance). The correlation functions are then calculated by smoothing using a moving average filter with a centered window of *±*100 microns. Error bars are *±*1 standard error of the mean within each window. Correlation functions (and the system-size scaling of correlations length) are qualitatively similar (though more or less noisy) for a range of window sizes from *±*25 up to *±*200 microns.

### Estimating System Size (*L*)

To estimate the size *L* of each population, we calculated the maximum pairwise separation (*L*(*t*) ≡ *d*_max_) between any two cells at each time point. The distance *L*(*t*) will fluctuate over time as the cells move. We define the size of the system as the mean value of this separation distance over the entire time series (*L* ≡ ⟨*L*(*t*) ⟩_*t*_). We found qualitatively similar results (i.e. scaling of correlation length with system size) if we alternatively defined the system size at each time point as *L*(*t*)= *d*_*con*_, where *d*_*con*_ = *A*^1*/*2^ and *A* is the area of the convex hull of all cell positions.

### Nonlocally coupled XY model as a data constrained maximum entropy model

To model collective motion in tumor cells, we used a maximum entropy framework. Maximum entropy models are required to match certain features of the data (in this case, the observed correlation *C*_int_, see below) but contain minimal additional statistical structure. Similar models have been previously used to describe firing patterns in neurons, immune system dynamics, interactions between antibiotics, and collective motion in starlings. Following [15], we constrain the model to match the scalar correlation over local neighborhoods of size *n*_*c*_, which is given by

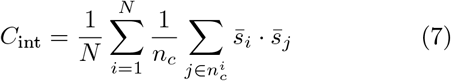

where *N* is the total number of cells, *n*_*c*_ is the integer size of the local neighborhood, and 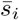 is the unit velocity vector describing the motion of cell *i*. The maximum entropy model consistent with this constraint is then given by [15]

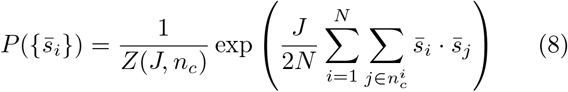

where 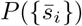 is the distribution over all configurations and *Z*(*J, n*_*c*_) is the partition function (i.e. the normalization constant). To fully specify the model, we must choose values of the parameters *J* and *n*_*c*_ such that the model reproduces the observed value of *C*_int_, a process that is equivalent to maximizing the likelihood that the model generates the configuration observed in a given snapshot of the flock.

To estimate model parameters, we first calculate the experimental correlation 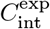 (where the superscript “exp” indicates this is the value observed in the experiment) for a single snapshot of the population. This experimental value must match the value of *C*_int_ produced by the model, which provides an explicit data-dependent constraint between the parameters *J* and *n*_*c*_,

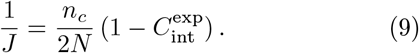

We then determine the remaining free parameter, *n*_*c*_, by numerically maximizing the log likelihood of the data given the model, which can be written as a function of the both 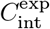 (a measured quantity) and *Z*(*J, n*_*c*_) (the partition function for the model).

Directly calculating the partition function *Z*(*J, n*_*c*_) is computationally expensive, even for this simple model. One option to simplify the calculation is to use a spin wave approximation, which provides an analytical expression for *Z*(*J, n*_*c*_) in terms of eigenvalues of a matrix *A* that describes the neighborhood structure of the population (i.e. which cells are in the local neighborhood, of size *n*_*c*_, of each cell). In the spin wave approximation, the partition function reduces to [15]

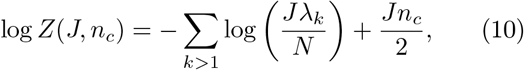

where *λ*_*k*_ are the eigenvalues of the matrix *A* = *δ*_*ij*_ Σ_*k*_ *n*_*ik*_ −*n*_*ij*_, where *n*_*ij*_ is 1 if cell *i* is in the local neighborhood of cell *j* and vice versa, 1/2 if cell *i* is in the local neighborhood of cell *j* but *j* is <u>not</u> in the local neighborhood of cell *i* (or vice versa), and 0 otherwise. It is important to note that the spin wave approximation is strictly valid only in highly polarized populations, as it relies on neglecting higher-order terms in an expansion velocity components perpendicular to the direction of polarization. In this work, we used the spin wave approximation to provide first-pass estimates of the parameters *J* and *n*_*c*_. While cell motion is not typically highly polarized in our dataset–and the spin wave approximation is therefore not strictly valid–we found that the parameters estimated from this approach do lead to strong agreement with the data.

### Estimating parameters from simulated data

To probe the reliability of parameters estimated with this spin wave approximation, we used Monte Carlo simulations to produce artificial data sets–snapshots of velocity configurations for populations of cells whose positions are identical to those in the experimental data–drawn from the model with specified values of *J* and *n*_*c*_. We then estimated the parameters directly from the simulated data using either 1) the spin wave approximation or 2) a direct least squares fitting to observed distribution *P* ({*θ*_*i*_}), where {*θ*_*i*_} is the configuration of velocity angles in the population.

### Mean field XY model

The maximum entropy model has two free parameters, *J* and *n*_*c*_, that are inferred from experimental data. In the limit *n*_*c*_ → *N* − 1 ≈ *N* (all-to-all coupling), this model reduces to a mean-field version of the 2D XY model with only a single free parameter, *J*. Estimating this parameter from data reduces to a classic inverse problem in statistical physics, with the log likelihood *L*(*J*) of the observed configurations taking the form

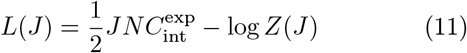

where 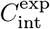 is now the average pairwise correlation taken over the entire population. Maximizing the likelihood is equivalent to requiring that the correlations measured experimentally 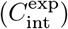 match those from the mean field model, which reduces to a classical inverse problem. In this case, we can write down an explicit expression for *J* in terms of the experimental measurable

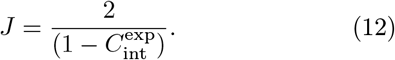

### Estimating critical parameters in systems with long-range coupling

Equation 12 highlights an important caveat of our approach when the range of coupling *n*_*c*_ is comparable to the total size of the system (*N*). The mean field XY model undergoes a phase transition at a critical value of *J*_*c*_ = 2. Equation 12 therefore indicates that observed data characterized by 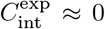 would appear to be at a critical point of the mean field model. In practice, of course, 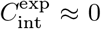 would also be expected for completely disordered populations–that is, for populations where the velocity direction is drawn from a uniform distribution. As a result, maximum likelihood estimates that indicate *J* ≈ *J*_*c*_ are not, alone, sufficient evidence of criticality, as they cannot distinguish systems in the disordered region (*J < J*_*c*_) from those at criticality. To quantitatively characterize these limitations for data sets comparable in size to our experimental data, we used Monte Carlo simulations to generate artificial data sets representing cell populations globally coupled with different values of *J*. We then calculated the maximum likelihood estimates of *J* from those in silico data sets (just as we did with experimental data). As expected, we found that estimates of *J* hover around the critical value *J*_*c*_ =2 for simulated systems at or below the critical point (Figure S7). On the other hand, estimates of *J* are accurate for systems slightly above the critical point.

### Monte Carlo Simulations and Phase Diagram

To characterize the phase diagram for the nonlocal XY model, we used Monte Carlo simulations to generate representative velocity distributions for different values of *n*_*c*_ and *J*. For each parameter pair, we calculated the mean polarization across trials and the generalized susceptibility χ, which corresponds (up to a proportionality constant) to the variance of the polarization across trials. We then estimated the critical surface to be the curve in parameter space that corresponds to a peak in χ. We also verified that the system exhibits sharply increasing polarization and a peak in the generalized heat capacity at the critical surface. For all simulations, cell positions (and therefore cell density) were taken from a representative experimental data set.

### Comparing velocity angle distributions between the model and data

To compare experimental results with results from the effective model, we calculated histograms of velocity angle. The model is symmetric under rotation and cannot provide information about the specific velocity direction– that is, we can globally rotate all velocities by an arbitrary angle. To compare histograms between the model and the data–or, for example, to combine data from multiple independent simulations (model) or multiple time points (experiment)–we first calculated the two frequency histograms to be compared. Then, we rotated all velocities in one population by an angle *θ*, and we tuned *θ* to achieve maximal alignment between the distributions (i.e. minimal difference in a least squares sense). Because the histograms themselves are noisy, this process could, in principle, lead to apparent similarities between distributions with fundamentally different shapes. Therefore, as a control, we simulated size-matched populations where velocity angles were drawn from a uniform distribution, and we then used this alignment process to compare these (nominally) uniform histograms with the histograms from the maximum entropy model. We confirmed that this alignment process does not yield substantial agreement between these different distributions (Figure S7).

## References

[1] D. N. Louis, A. Perry, P. Wesseling, D. J. Brat, I. A. Cree, D. Figarella-Branger, C. Hawkins, H. K. Ng, S. M. Pfister, G. Reifenberger, R. Soffietti, A. von Deimling, and D. W. Ellison, Neuro Oncol 23, 1231 (2021).

[2] M. Ceccarelli, F. P. Barthel, T. M. Malta, T. S. Sabedot, S. R. Salama, B. A. Murray, O. Morozova, Y. Newton, A. Radenbaugh, S. M. Pagnotta, S. Anjum, J. Wang, G. Manyam, P. Zoppoli, S. Ling, A. A. Rao, M. Grifford, A. D. Cherniack, H. Zhang, L. Poisson, J. Carlotti, C. G., D. P. Tirapelli, A. Rao, T. Mikkelsen, C. C. Lau, W. K. Yung, R. Rabadan, J. Huse, D. J. Brat, N. L. Lehman, J. S. Barnholtz-Sloan, S. Zheng, K. Hess, G. Rao, M. Meyerson, R. Beroukhim, L. Cooper, R. Akbani, M. Wrensch, D. Haussler, K. D. Aldape, P. W. Laird, D. H. Gutmann, H. Noushmehr, A. Iavarone, and R. G. Verhaak, Cell 164, 550 (2016).

[3] R. B. Puchalski, N. Shah, J. Miller, R. Dalley, S. R. Nomura, J. G. Yoon, K. A. Smith, M. Lankerovich, D. Bertagnolli, K. Bickley, A. F. Boe, K. Brouner, S. Butler, S. Caldejon, M. Chapin, S. Datta, N. Dee, T. Desta, T. Dolbeare, N. Dotson, A. Ebbert, D. Feng, X. Feng, M. Fisher, G. Gee, J. Goldy, L. Gourley, B. W. Gregor, G. Gu, N. Hejazinia, J. Hohmann, P. Hothi, R. Howard, K. Joines, A. Kriedberg, L. Kuan, C. Lau, F. Lee, H. Lee, T. Lemon, F. Long, N. Mastan, E. Mott, C. Murthy, K. Ngo, E. Olson, M. Reding, Z. Riley, D. Rosen, D. Sandman, N. Shapovalova, C. R. Slaughterbeck, A. Sodt, G. Stockdale, A. Szafer, W. Wakeman, P. E. Wohnoutka, S. J. White, D. Marsh, R. C. Rostomily, L. Ng, C. Dang, A. Jones, B. Keogh, H. R. Gittleman, J. S. Barnholtz-Sloan, P. J. Cimino, M. S. Uppin, C. D. Keene, F. R. Farrokhi, J. D. Lathia, M. E. Berens, A. Iavarone, A. Bernard, E. Lein, J. W. Phillips, S. W. Rostad, C. Cobbs, M. J. Hawrylycz, and G. D. Foltz, Science 360, 660 (2018).

[4] T. Daubon, R. Magaut, and A. Bikfalvi, Adv Exp Med Biol 1329, 109 (2021).

[5] A. Vollmann-Zwerenz, V. Leidgens, G. Feliciello, C. A. Klein, and P. Hau, Int J Mol Sci 21 (2020), 10.3390/ijms21061932.

[6] P. G. Gritsenko, O. Ilina, and P. Friedl, J Pathol 226, 185 (2012).

[7] A. Comba, S. Motsch, P. J. Dunn, T. C. Hollon, A. E. Argento, D. B. Zamler, P. E. Kish, A. Kahana, C. G. Kleer, M. G. Castro, and P. R. Lowenstein, Nature Communication 13, 3606 (2022).

[8] M. Alieva, V. Leidgens, M. J. Riemenschneider, C. A. Klein, P. Hau, and J. van Rheenen, Sci Rep 9, 2054 (2019).

[9] A. Haeger, M. Krause, K. Wolf, and P. Friedl, Biochim Biophys Acta 1840, 2386 (2014).

[10] S. Jamous, A. Comba, P. R. Lowenstein, and S. Motsch, PLoS computational biology 16, e1007611 (2020).

[11] P. G. Gritsenko, N. Atlasy, C. E. J. Dieteren, A. C. Navis, J. H. Venhuizen, C. Veelken, D. Schubert, A. AckerPalmer, B. A. Westerman, T. Wurdinger, W. Leenders, P. Wesseling, H. G. Stunnenberg, and P. Friedl, Nat Cell Biol 22, 97 (2020).

[12] C. J. Liu, G. A. Shamsan, T. Akkin, and D. J. Odde, Biophys J 117, 1179 (2019).

[13] A. Cavagna, A. Cimarelli, I. Giardina, G. Parisi, R. Santagati, F. Stefanini, and M. Viale, Proceedings of the National Academy of Sciences 107, 11865 (2010).

[14] G. F. Young, L. Scardovi, A. Cavagna, I. Giardina, and N. E. Leonard, PLoS computational biology 9 (2013).

[15] W. Bialek, A. Cavagna, I. Giardina, T. Mora, E. Silvestri, M. Viale, and A. M. Walczak, Proceedings of the National Academy of Sciences 109, 4786 (2012).

[16] W. Bialek, A. Cavagna, I. Giardina, T. Mora, O. Pohl, E. Silvestri, M. Viale, and A. M. Walczak, Proceedings of the National Academy of Sciences 111, 7212 (2014).

[17] T. Niizato, H. Murakami, and Y.-P. Gunji, Biosystems 119, 62 (2014).

[18] H. Ling, G. E. Mclvor, K. van der Vaart, R. T. Vaughan, A. Thornton, and N. T. Ouellette, Proceedings of the Royal Society B 286, 20190865 (2019).

[19] U. Lopez, J. Gautrais, I. D. Couzin, and G. Theraulaz, Interface focus 2, 693 (2012).

[20] C. Hemelrijk, D. Reid, H. Hildenbrandt, and J. Padding, Fish and Fisheries 16, 511 (2015).

[21] A. Sokolov and I. S. Aranson, Physical review letters 109, 248109 (2012).

[22] H.-P. Zhang, A. Be’er, E.-L. Florin, and H. L. Swinney, Proceedings of the National Academy of Sciences 107, 13626 (2010).

[23] H. Wioland, E. Lushi, and R. E. Goldstein, New Journal of Physics 18, 075002 (2016).

[24] O. Feinerman, I. Pinkoviezky, A. Gelblum, E. Fonio, and N. S. Gov, Nature Physics 14, 683 (2018).

[25] J. Buhl, D. J. Sumpter, I. D. Couzin, J. J. Hale, E. Despland, E. R. Miller, and S. J. Simpson, Science 312, 1402 (2006).

[26] A. Attanasi, A. Cavagna, L. Del Castello, I. Giardina, S. Melillo, L. Parisi, O. Pohl, B. Rossaro, E. Shen, E. Silvestri, et al., PLoS computational biology 10, e1003697 (2014).

[27] N. Ouellette, Physical Biology (2022).

[28] K. van der Vaart, M. Sinhuber, A. M. Reynolds, and N. T. Ouellette, Journal of The Royal Society Interface 17, 20200018 (2020).

[29] N. Bain and D. Bartolo, Science 363, 46 (2019).

[30] F. Ginelli, F. Peruani, M.-H. Pillot, H. Chaté, G. Theraulaz, and R. Bon, Proceedings of the National Academy of Sciences 112, 12729 (2015).

[31] A. Bottinelli, D. T. Sumpter, and J. L. Silverberg, Physical review letters 117, 228301 (2016).

[32] A. Czirók, A.-L. Barabási, and T. Vicsek, Physical Review Letters 82, 209 (1999).

[33] A. Cavagna and I. Giardina, Behav Processes 84, 653 (2010).

[34] T. Vicsek and A. Zafeiris, Physics reports 517, 71 (2012).

[35] E. Méhes and T. Vicsek, Integrative biology 6, 831 (2014).

[36] N. Shimoyama, K. Sugawara, T. Mizuguchi, Y. Hayakawa, and M. Sano, Physical Review Letters 76, 3870 (1996).

[37] A. V. Savkin, IEEE Transactions on Automatic Control 49, 981 (2004).

[38] G. Vásárhelyi, C. Virágh, G. Somorjai, T. Nepusz, A. E. Eiben, and T. Vicsek, Science Robotics 3, eaat3536 (2018).

[39] L. Qin, D. Yang, W. Yi, H. Cao, and G. Xiao, Mol Biol Cell 32, 1267 (2021).

[40] J. Zhang, K. F. Goliwas, W. Wang, P. V. Taufalele, F. Bordeleau, and C. A. Reinhart-King, Proc Natl Acad Sci U S A 116, 7867 (2019).

[41] E. Cristiani, M. Menci, M. Papi, and L. Brafman, J Math Biol 83, 45 (2021).

[42] H. Trenchard and M. Perc, Biosystems 147, 40 (2016).

[43] K. Sonoda, H. Murakami, T. Niizato, T. Tomaru, Y. Nishiyama, and Y. P. Gunji, Biosystems 185, 104019 (2019).

[44] T. Mora and W. Bialek, Journal of Statistical Physics 144, 268 (2011).

[45] P. Bak and M. Paczuski, Proceedings of the National Academy of Sciences 92, 6689 (1995).

[46] P. M. Chaikin, T. C. Lubensky, and T. A. Witten, Principles of condensed matter physics, vol. 10 (Cambridge university press Cambridge, 1995).

[47] M. A. Munoz, Reviews of Modern Physics 90, 031001 (2018).

[48] G. Tkačik, T. Mora, O. Marre, D. Amodei, S. E. Palmer, M. J. Berry, and W. Bialek, Proceedings of the National Academy of Sciences 112, 11508 (2015).

[49] M. Rubinov, O. Sporns, J.-P. Thivierge, and M. Breakspear, PLoS computational biology 7, e1002038 (2011).

[50] D. Fraiman and D. R. Chialvo, Front. Physiol. 3, 307 (2012).

[51] A. Haimovici, E. Tagliazucchi, P. Balenzuela, and D. R. Chialvo, Phys. Rev. Lett. 110, 178101 (2013).

[52] P. Rämö, J. Kesseli, and O. Yli-Harja, Journal of Theoretical Biology 242, 164 (2006).

[53] B. C. Daniels, H. Kim, D. Moore, S. Zhou, H. B. Smith, B. Karas, S. A. Kauffman, and S. I. Walker, Physical review letters 121, 138102 (2018).

[54] Q.-Y. Tang, Y.-Y. Zhang, J. Wang, W. Wang, and D. R. Chialvo, Physical review letters 118, 088102 (2017).

[55] A. Attanasi, A. Cavagna, L. Del Castello, I. Giardina, S. Melillo, L. Parisi, O. Pohl, B. Rossaro, E. Shen, E. Silvestri, et al., Physical review letters 113, 238102 (2014).

[56] A. Comba, P. J. Dunn, A. E. Argento, P. Kadiyala, M. Ventosa, P. Patel, D. B. Zamler, F. J. Nunez, L. Zhao, and M. G. Castro, Neuro-oncology 22, 806 (2020).

[57] F. J. Núñez, F. M. Mendez, P. Kadiyala, M. S. Alghamri, M. G. Savelieff, M. B. Garcia-Fabiani, S. Haase, C. Koschmann, A.-A. Calinescu, N. Kamran, M. Saxena, R. Patel, S. Carney, M. Z. Guo, M. Edwards, M. Ljungman, T. Qin, M. A. Sartor, R. Tagett, S. Venneti, J. Brosnan-Cashman, A. Meeker, V. Gorbunova, L. Zhao, D. M. Kremer, L. Zhang, C. A. Lyssiotis, L. Jones, C. J. Herting, J. L. Ross, D. Hambardzumyan, S. Hervey-Jumper, M. E. Figueroa, P. R. Lowenstein, and M. G. Castro, 11, eaaq1427 (2019).

[58] C. Koschmann, A. A. Calinescu, F. J. Nunez, A. Mackay, J. Fazal-Salom, D. Thomas, F. Mendez, N. Kamran, M. Dzaman, L. Mulpuri, J. Krasinkiewicz, R. Doherty, R. Lemons, J. A. Brosnan-Cashman, Y. Li, S. Roh, L. Zhao, H. Appelman, D. Ferguson, V. Gorbunova, A. Meeker, C. Jones, P. R. Lowenstein, and M. G. Castro, Sci Transl Med 8, 328ra28 (2016).

[59] E. Schneidman, M. J. Berry, R. Segev, and W. Bialek, Nature 440, 1007 (2006).

[60] J. Shlens, G. D. Field, J. L. Gauthier, M. I. Grivich, D. Petrusca, A. Sher, A. M. Litke, and E. Chichilnisky, Journal of Neuroscience 26, 8254 (2006).

[61] A. Cavagna, I. Giardina, and T. S. Grigera, Physics Reports The Physics of Flocking: Correlation as a Compass from Experiments to Theory, 728, 1 (2018).

[62] J. N. Kapur, Maximum-entropy models in science and engineering (John Wiley & Sons, 1989).

[63] K. Wood, S. Nishida, E. D. Sontag, and P. Cluzel, Proceedings of the National Academy of Sciences 109, 12254 (2012).

[64] G. Yeo and C. B. Burge, Journal of computational biology 11, 377 (2004).

[65] E. T. Jaynes, Physical review 106, 620 (1957).

[66] H. C. Nguyen, R. Zecchina, and J. Berg, Advances in Physics 66, 197 (2017).

[67] M. Castellana and W. Bialek, Physical review letters 113, 117204 (2014).

[68] R. R. Stein, D. S. Marks, and C. Sander, PLoS computational biology 11, e1004182 (2015).

[69] L. R. Mead and N. Papanicolaou, Journal of Mathematical Physics 25, 2404 (1984).

[70] J. Hidalgo, J. Grilli, S. Suweis, M. A. Munoz, J. R. Banavar, and A. Maritan, Proceedings of the National Academy of Sciences 111, 10095 (2014).

[71] J. Humplik and G. Tkačik, PLoS computational biology 13, e1005763 (2017).

[72] N. Stollenwerk and V. Jansen, Population Biology and Criticality (World Scientific, 2011).

[73] M. Osswald, E. Jung, F. Sahm, G. Solecki, V. Venkataramani, J. Blaes, S. Weil, H. Horstmann, B. Wiestler, M. Syed, et al., Nature 528, 93 (2015).

[74] T. Mora, A. M. Walczak, L. Del Castello, F. Ginelli, S. Melillo, L. Parisi, M. Viale, A. Cavagna, and I. Giardina, Nature physics 12, 1153 (2016).

[75] A. Cavagna, I. Giardina, F. Ginelli, T. Mora, D. Piovani, R. Tavarone, and A. M. Walczak, Physical Review E 89, 042707 (2014).

[76] Y. Roudi, S. Nirenberg, and P. E. Latham, PLoS computational biology 5, e1000380 (2009).

[77] L. Merchan and I. Nemenman, Journal of Statistical Physics 162, 1294 (2016).

[78] I. Mastromatteo and M. Marsili, Journal of Statistical Mechanics: Theory and Experiment 2011, P10012 (2011).

[79] D. J. Schwab, I. Nemenman, and P. Mehta, Physical review letters 113, 068102 (2014).

[80] M. Nonnenmacher, C. Behrens, P. Berens, M. Bethge, and J. H. Macke, PLoS computational biology 13, e1005718 (2017).

[81] R. J. Cubero, J. Jo, M. Marsili, Y. Roudi, and J. Song, Journal of Statistical Mechanics: Theory and Experiment 2019, 063402 (2019).

[82] R. Ni and N. Ouellette, The European Physical Journal Special Topics 224, 3271 (2015).

[83] D. A. Martin, T. L. Ribeiro, S. A. Cannas, T. S. Grigera, D. Plenz, and D. R. Chialvo, Scientific Reports 11, 1 (2021).

