## Supplemental Text for "Scale-free correlations and criticality in an experimental model of brain cancer"

### SUPPLEMENTARY INFORMATION

#### Estimating parameters for maximum entropy and mean field models is insufficient to distinguish between critical and random (sub-critical) systems.

To explore the limits of our approach for gauging proximity to a critical point in experimental populations, we used Monte Carlo simulations to generate artificial data sets generated by either 1) the maximum entropy model or 2) the mean field model across a range of parameter values. We then estimated the best-fit values of $J$ and $n_c$ (for the full model) or $J$ alone (for the mean field model) and compared those to the "true" values used in the simulations (Figure S8 and Figure S9). In both models, parameter combinations in the super-critical (ordered) regime lead to parameter estimates that are also super-critical, though the approach systematically over-estimates $J$, meaning some systems just above the critical point will artificially appear to be farther from criticality. On the other hand, parameter estimates for systems that are in the sub-critical regime–those that fall below the critical curve in the full model, or those characterized by $J < J_c = 2$ in the mean field model–fall near the critical point, even when the system itself is subcritical.

This analysis underscores the need for caution when inferring criticality using only experimentally derived estimates of model parameters. While this approach is sufficient to distinguish data sets far above the critical point–that is, populations with relatively large degrees of order–from those near the critical point, it cannot, alone, distinguish between populations poised near criticality and those (even far below) the critical point. Nevertheless, such models–in combination with model-independent metrics (e.g. scale-free correlations)–may provide substantial evidence for criticality. Indeed, in the case of glioma populations, we find evidence of non-random ordering in the form of scale-free correlations that would not be expected in highly disordered populations far below the critical point. At the same time, the maximum entropy analysis suggests that these populations are not poised far above the critical point–that is, they do not appear consistent with fully ordered populations. Taken together, these results suggest that glioma populations are poised near a critical point–just on the edge between the ordered and disordered state.

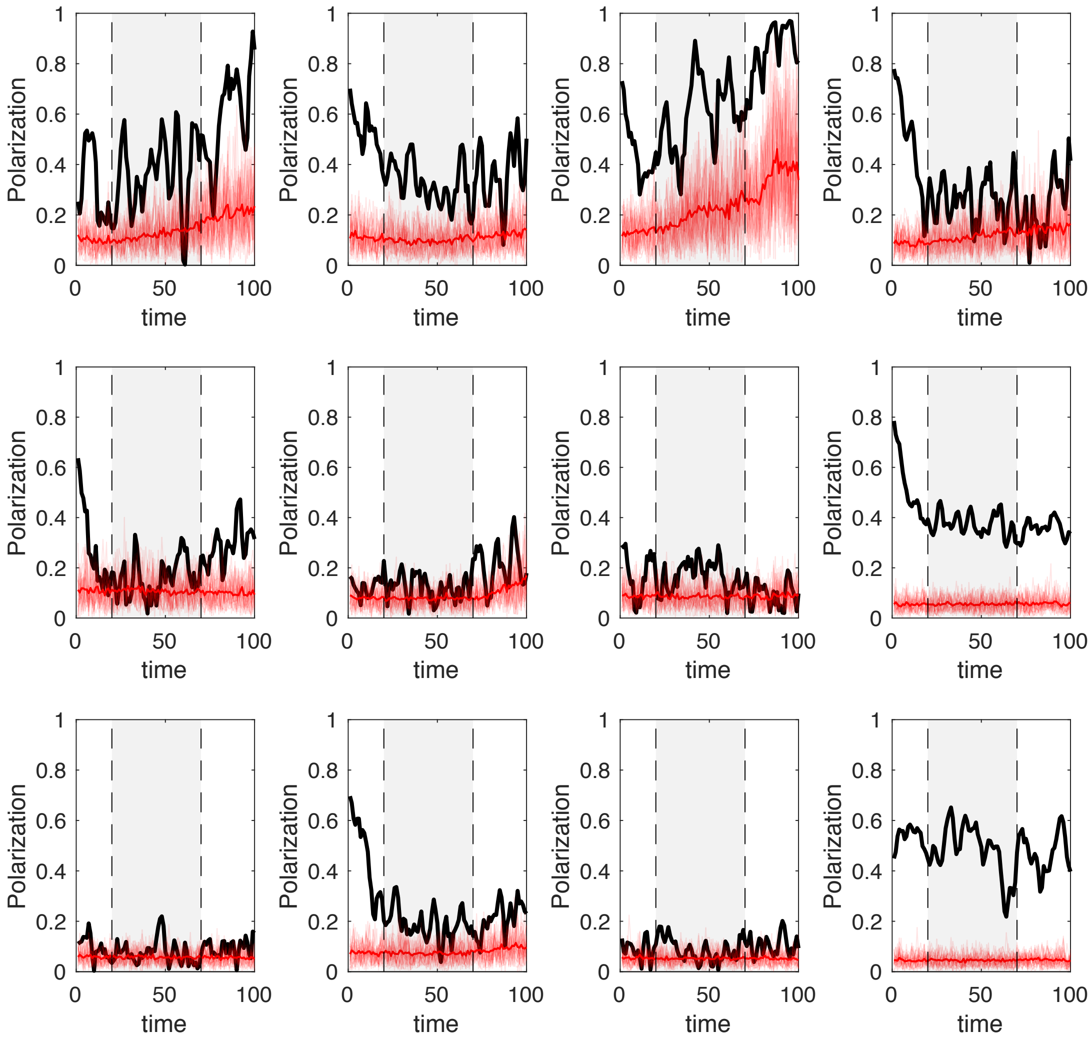


FIG. S1. **Polarization over time for all populations.** Polarization over time for all 12 populations (black) and size-matched simulated populations with velocities drawn from uniform distribution on the circle (red). Gray shaded region corresponds to time period where analyses were performed. This period was chosen to 1) allow initial transient behavior to subside and 2) minimize effects of later time points ($t > 70$) when cell numbers decline rapidly as cells escape field of view.

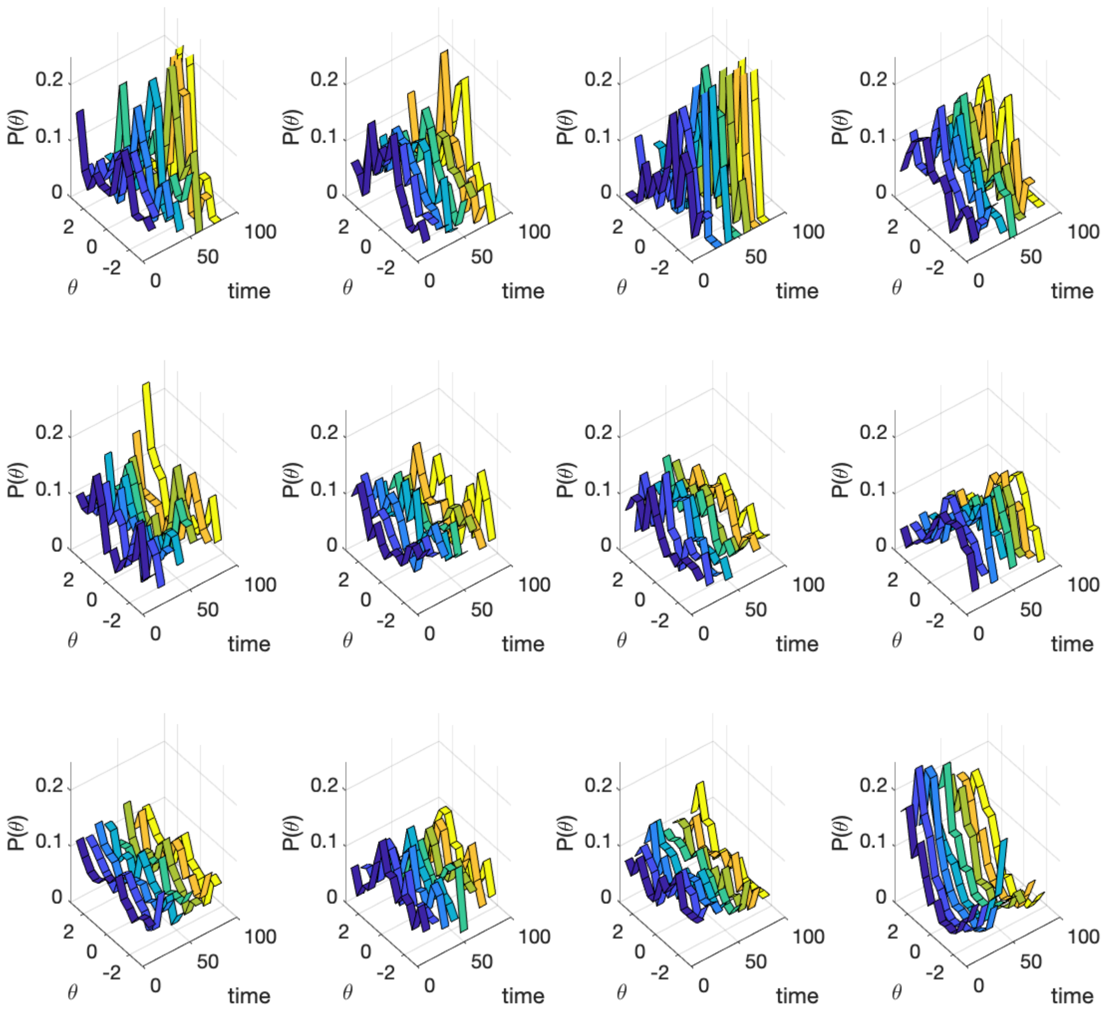


FIG. S2. **Velocity angle histograms over time.** Histograms of velocity angle over all cells in a population over time.

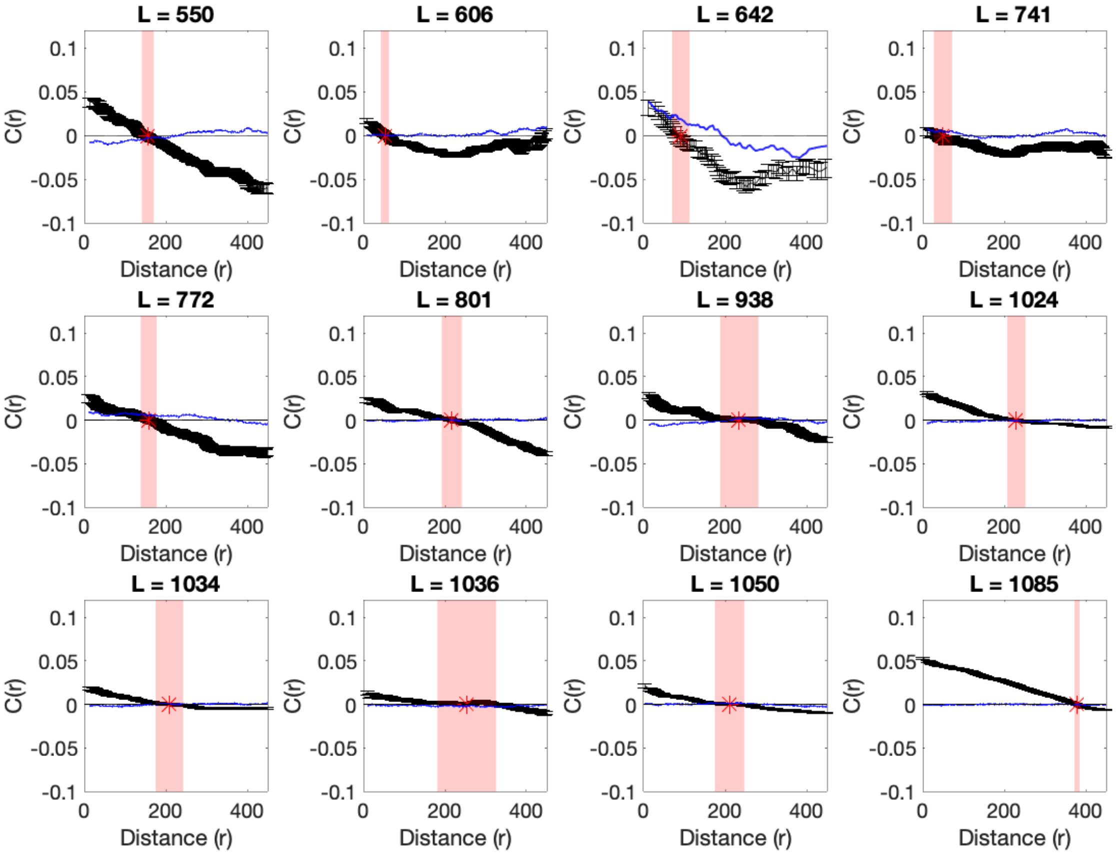


FIG. S3. **Correlation functions for populations of different sizes.** Correlation functions illustrate correlations between directional velocity for cells separated by a given distance (r; in microns) for populations with different sizes $L$. Correlation functions are defined by $C(r) = \langle \delta s_i \cdot \delta s_j \rangle_r$, where $\delta s_i \equiv s_i - \langle s \rangle$ is the velocity of cell $i$ in the moving reference frame where the population average velocity ($\langle s \rangle$) has been subtracted out. Angle brackets $\langle \rangle_r$ indicate an average taken over all cells separated by distance $r$. Black markers: time-averaged correlations in a given population; red shaded region indicates estimated correlation length $\xi$, which corresponds to the crossover point $C(\xi) = 0$. Blue curves are from simulations of size-matched populations where velocity angle for each cell is chosen from a uniform distribution on the circle.

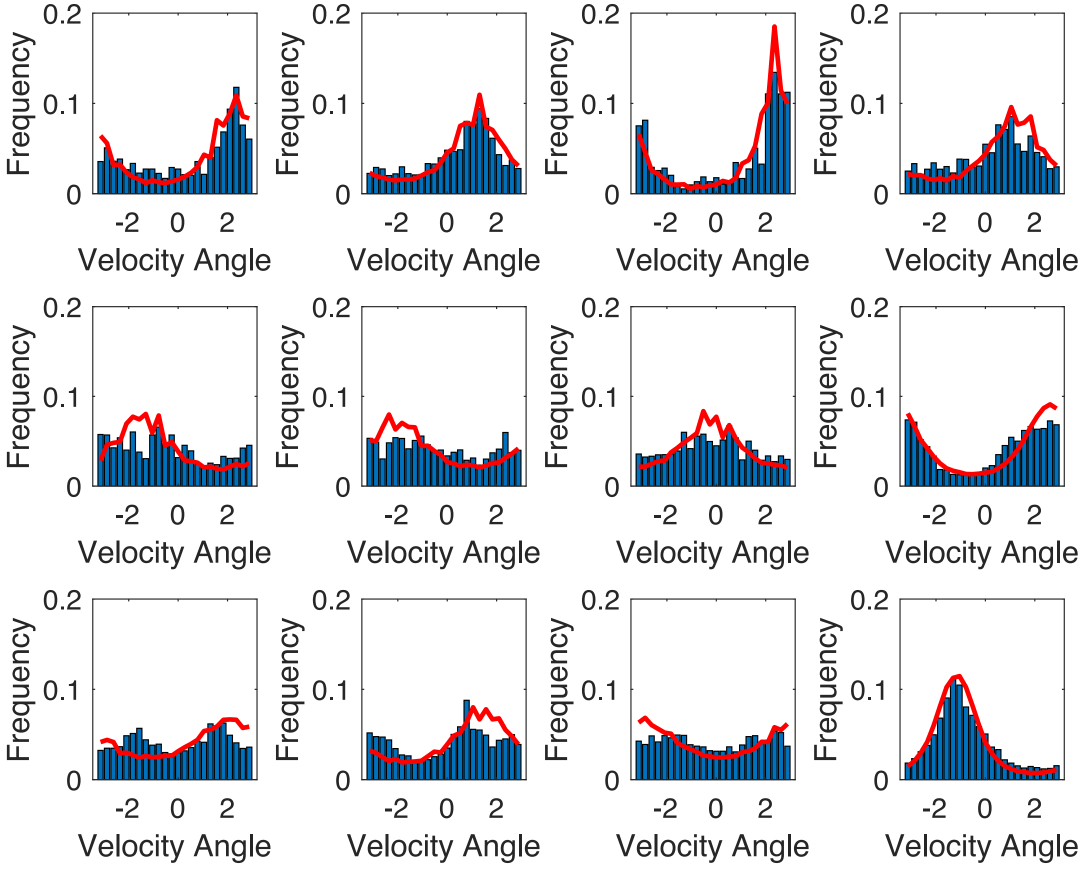


FIG. S4. **Maximum entropy model approximately captures distribution of velocity angles.** Histograms of velocity direction (angle) for experiment (blue) and the maximum entropy model (red) for the same populations as in Figure S3.

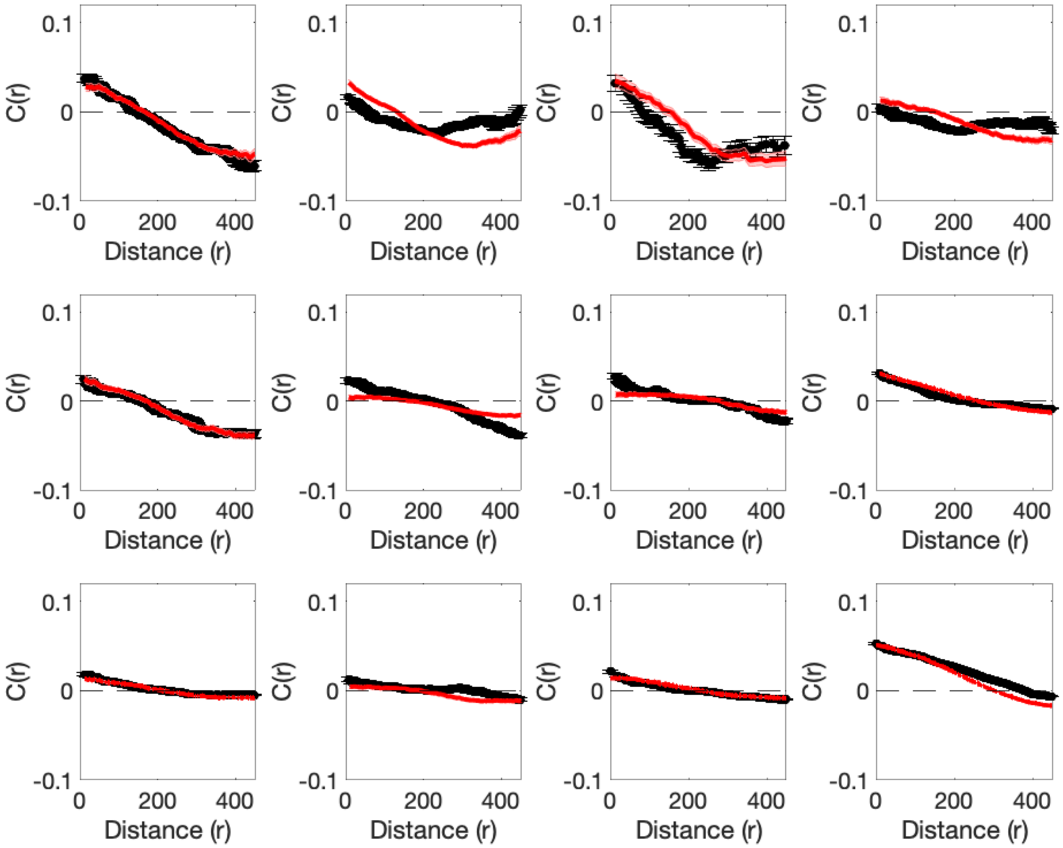


FIG. S5. **Maximum entropy model captures most qualitative features of correlation functions.** Time averaged correlation functions $C(r)$ for experimental data (black) and the maximum entropy model (red) with best-fit parameters $n_c$ and $J$. The correlation function is defined by $C(r) = \langle \delta s_i \cdot \delta s_j \rangle_r$, where $\delta s_i \equiv s_i - \langle s \rangle$ is the velocity of cell $i$ in the moving reference frame where the population average velocity ($\langle s \rangle$) has been subtracted out. Angle brackets $\langle \rangle_r$ indicate an average taken over all cells separated by distance $r$.

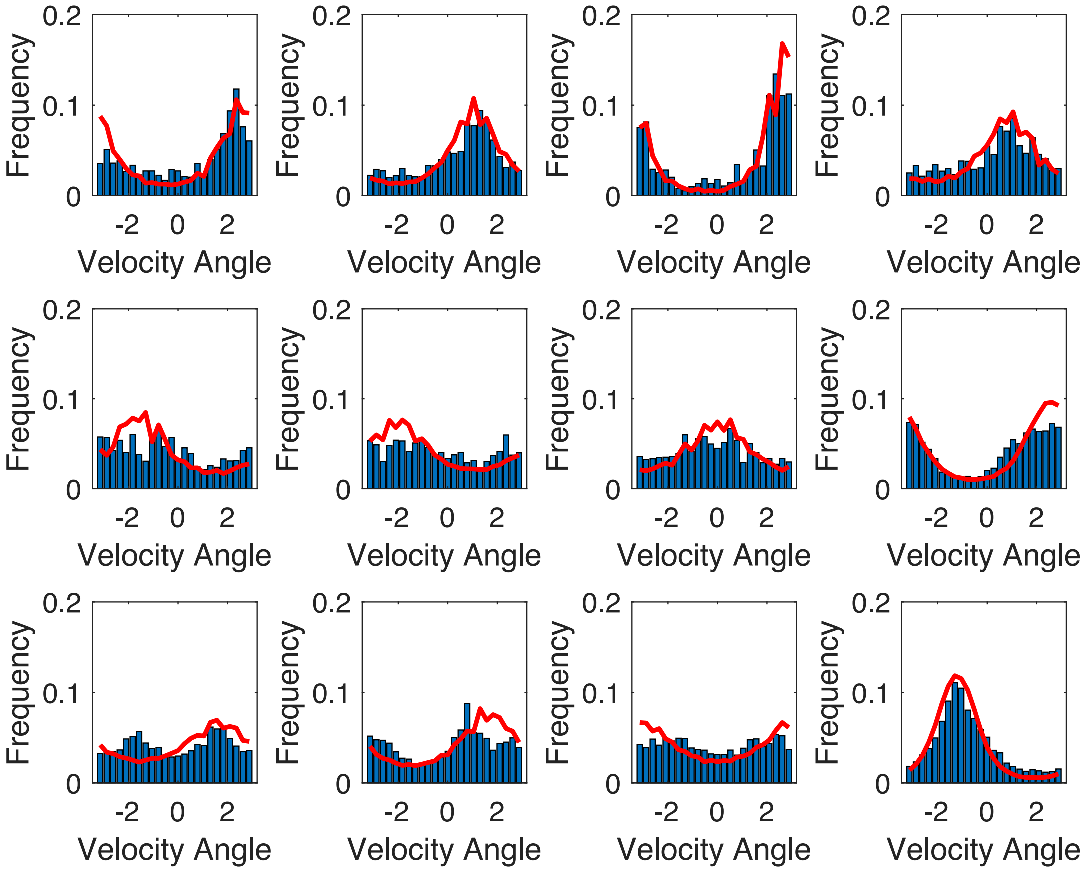


FIG. S6. **Mean field model XY model approximately captures distribution of velocity angles.** Histograms of velocity direction (angle) for experiment (blue) and the mean field XY model (red) for the same populations as in Figure S3.

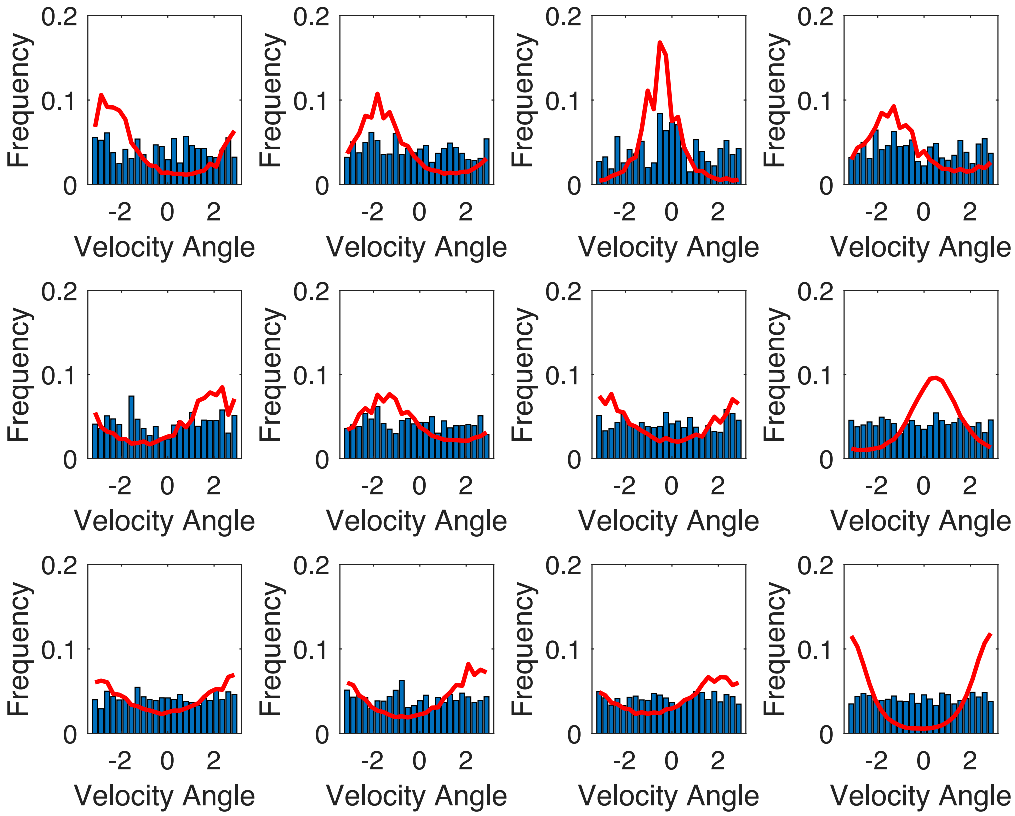


FIG. S7. **Histograms of angular velocities estimated from model differ from those of size-matched populations with random velocity orientations.** Histograms of velocity direction (angle) for the mean field XY model (red) and those for population-size matched populations where velocity angles were randomly drawn from a uniform distribution on the circle.

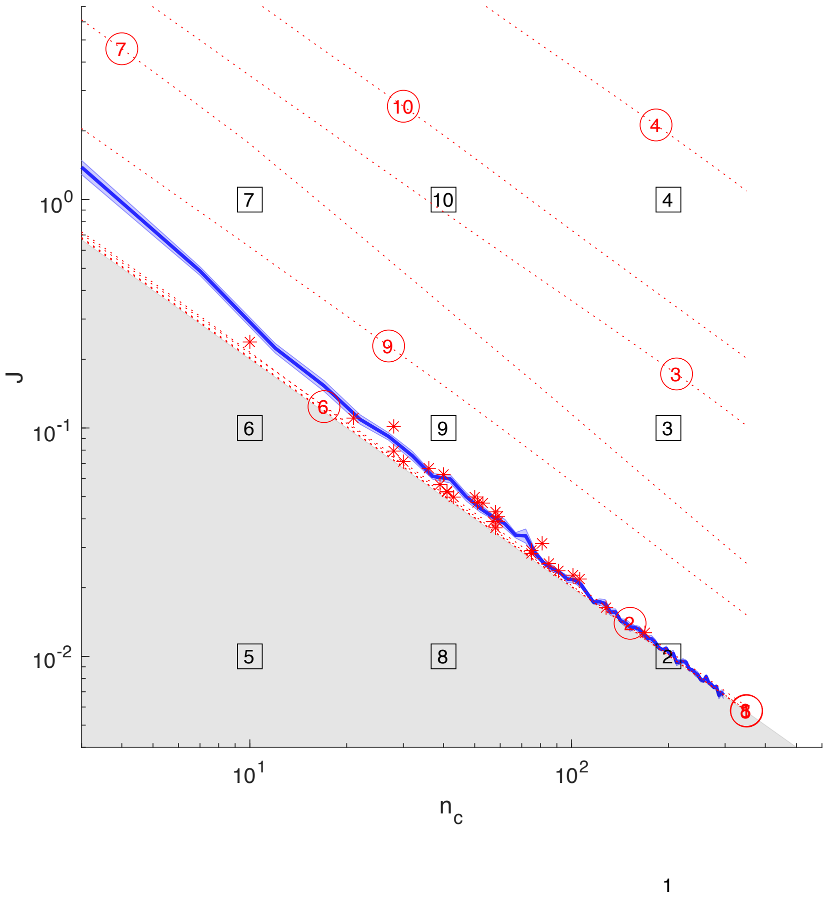


FIG. S8. **Estimating parameters from simulated data sets.** Phase diagram with phase boundary (blue line; shaded region indicates $\pm$ standard error) estimated from Monte Carlo simulations. Gray region indicates parameter regime ($J < 2/n_c$) forbidden when empirical data exhibits positive semidefinite correlations $C_{int} \geq 0$. Numbers indicate true parameter values (squares) and estimated parameter values (circles) for simulated data sets. Red stars indicate estimated parameters for experimental data from Figure 1.

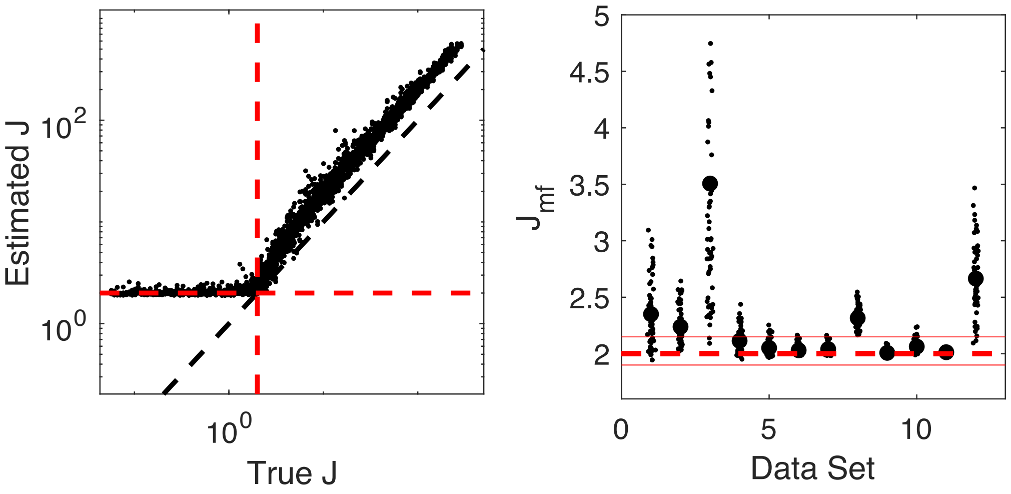


FIG. S9. **Mean field model illustrates limitation of approach in sub-critical regions** Left panel: to evaluate the estimation procedure for the mean field model, we simulated an all-to-all connected model over a range of coupling ($J$) values and then estimated the coupling values from snapshots of the model (i.e. collections of position and velocity vectors drawn from a Monte Carlo simulation). We found that below the critical point ($J_c = 2$) the estimated value of $J$ (vertical axis) is always near the critical value, indicating that sub-critical systems cannot be distinguished from those at or near the critical point. Right panel: For each of the twelve data sets in this paper, we estimated the coupling parameter ($J_{mf}$) in a model that assumes all-to-all coupling (so $n_c \rightarrow N-1$). Small points: individual snapshots over time; large points: mean over all time points. While in some cases (e.g. data sets 8, 12) the estimated value of $J_{mf}$ is clearly larger than the critical value $J_c = 2$, in many other cases the estimated values are very near the critical value, and this analysis along cannot distinguish between criticality and sub-criticality.

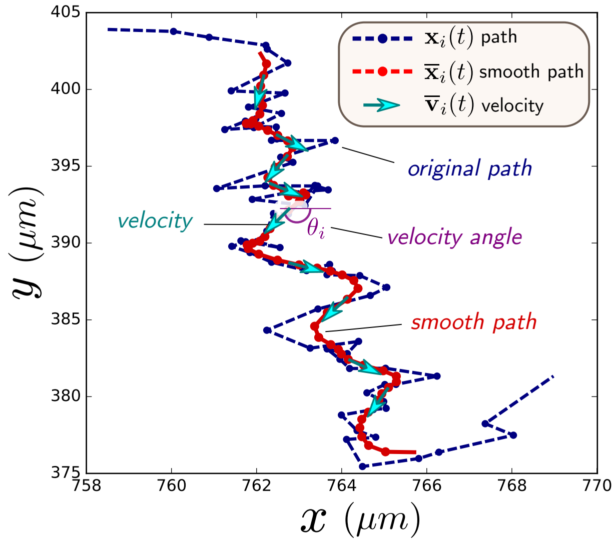


FIG. S10. Example of path obtained with ImageJ (blue dashed line) and the filtered path (red) obtained after filtering. The smooth path allows to estimate the velocity of the cell at each time step.

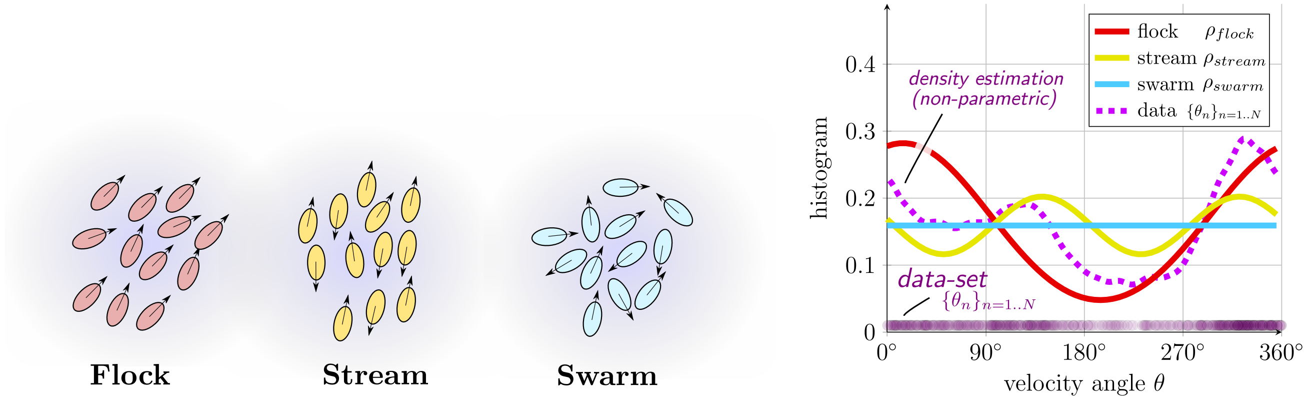


FIG. S11. Classification of pattern formations. **Left**: Sketch configuration of the three patterns identified: flock, stream and swarm. **Right**: corresponding velocity angles distribution for the three patterns.
